# Transmission Expression Signature in Nascent *Plasmodium vivax* Blood Stage Infection

**DOI:** 10.1101/175018

**Authors:** Swamy Rakesh Adapa, Rachel A. Taylor, Chengqi Wang, Richard Thomson-Luque, Leah R. Johnson, Rays H.Y. Jiang

**Affiliations:** Department of Global Health (GH) & Center for Drug Discovery and Innovation (CDDI), College of Public Health, University of South Florida. Tampa, FL 33612, USA; Department of Integrative Biology, University of South Florida. Tampa, FL, USA

## Abstract

The lack of a continuous *in vitro* culture system for *Plasmodium vivax* severely limits our knowledge of pathophysiology of the most widespread malaria parasite. To gain direct understanding of *P. vivax* human infections, we used Next Generation Sequencing data mining to unravel parasite *in vivo* expression profiles for *P. vivax,* and *P. falciparum* as comparison. We performed cloud and local computing to extract parasite transcriptomes from publicly available raw data of human blood samples. We developed a Poisson Modelling (PM) method to confidently identify parasite derived transcripts in mixed RNAseq signals of infected host tissues. We successfully retrieved and reconstructed parasite transcriptomes from infected patient blood as early as the first blood stage cycle; and the same methodology did not recover any significant signal from controls. Surprisingly, these first generation blood parasites already show strong signature of transmission, which indicates the commitment from asexual-to-sexual stages. Further, we develop mathematical models for *P. vivax* and *P. falciparum* to assess the epidemiological impact of possible 7-day early stage transmission and *P. vivax* complex life cycle. The study uncovers the earliest onset of *P. vivax* blood pathogenesis and highlights the challenges of *P. vivax* eradication programs.

**Author summary:** We discovered that *P. vivax in vivo* parasitemia is associated with gametocytogenesis expression signature within the first blood stage cycle, that is, eight days from a mosquito bite. Our results suggest that asexual-to-sexual commitment may happen with first generation merozoite infection. This allows for the possibility of transmission at this early stage, much earlier than for *P. falciparum*. Our novel mathematical model accounts for multiple unique aspects of *P. vivax* biology to advance our understanding of expected disease prevalence, and compares the results to those of *P. falciparum*. We demonstrate that given the presence of asymptotical carriers and the possibility of relapses, earlier parasite transmission is capable of increasing the spread of disease within human populations. In summary, *P. vivax* gametogenesis has the potential to fast track the transmission cycle, which will drive enhanced propagation of the disease during the transmission season and clinical relapses.

## Introduction

*Plasmodium vivax (P. vivax)* infection has the most widespread distribution across different continents of any malaria parasite, with up to 2.6 billion people estimated to be at risk [1]. It can lead to severe disease and death but, despite the high disease burden [2], there is a lack of in-depth understanding of the distinct pathogenesis of *P. vivax*. This has resulted in a lack of targeted control measures. Thus, as malaria cases decline overall, the proportion of cases attributable to *P. vivax* is on the rise [3].

*P. vivax* has a complex transmission cycle with distinct biological features compared to other malaria parasites, most notably: the high prevalence of asymptomatic carriers and the potential for disease relapses; and gametocytes in circulation at the very beginning of infections. In contrast to the better studied *Plasmodium falciparum*, *P. vivax* has the unique ability to remain as dormant hypnozoites in a hepatocyte in the liver and, in the future, to reactivate a blood stage infection leading to what is termed a clinical relapse [4, 5]. Unlike *P. falciparum*, there are currently no established laboratory methods to culture *P. vivax in vitro* [6]. Furthermore, in *P. vivax*, the merozoites from both exo-erythrocytic and intra-erythrocytic schizogony only successfully infect reticulocytes [6], which typically comprise about one percent of red blood cell. This leads to low parasitemia rates in peripheral circulation. The host requirement of human reticulocytes and many other technical challenges hampers studies of this parasite. These unique *P. vivax* life cycle characteristics pose major challenges for the understanding of *P. vivax* pathogenesis and hence the elimination of malaria worldwide [5].

Human malaria infection starts with the inoculation of sporozoites into the skin dermis through the proboscis of female *Anopheles* mosquitoes, where it is hosted in her salivary glands. Some part of the inoculum enters the bloodstream and within a few minutes they invade hepatocytes in the liver [7, 8]. During the next five to eight days (depending on the Plasmodium spp), the parasite transforms into a large exoerythrocytic form, packed with thousands of merozoites inside a parasitophorous vacuolar membrane (PVM). As the parasite matures the membrane breaks down into small packets of vesicles filled with merozoites. These are released into the blood stream, leading to erythrocytic invasion [9]. In the next 48 hours the parasite undergoes mitotic division and cytoplasmic growth inside the erythrocyte. They may develop either directly into a schizont (asexual) or gametocyte (sexual) [5]. For *P. falciparum* the sexual stages are not found in the periphery until after multiple blood stage cycles because gametogenesis, which requires bone marrow sequestration, takes 10 to 12 days, to achieve the fully transmissible stage V gametocyte [10]. In contrast, the appearance of *P. vivax* sexual stages is believed to be much earlier [5]. However, whether sexual commitment in *P. vivax* occurs early still needs to be determined.

In this study (Fig 1), we directly examine patient blood sequencing data to recover *P. vivax* transcripts in the earliest time point possible during the blood stage, i.e. immediately after sporozoite invasion and the liver stage parasite ruptures into the blood stream. We discovered a very early gametogenesis expression signature, indicating the possibility of very early sexual commitment and possible transmission. Lastly, to evaluate the epidemiological impact of this possible early transmission, we constructed a mathematical model of *P. vivax* transmission, which quantifies the effect of relapses, asymptomatic carriers and early transmission.

**Fig 1.**
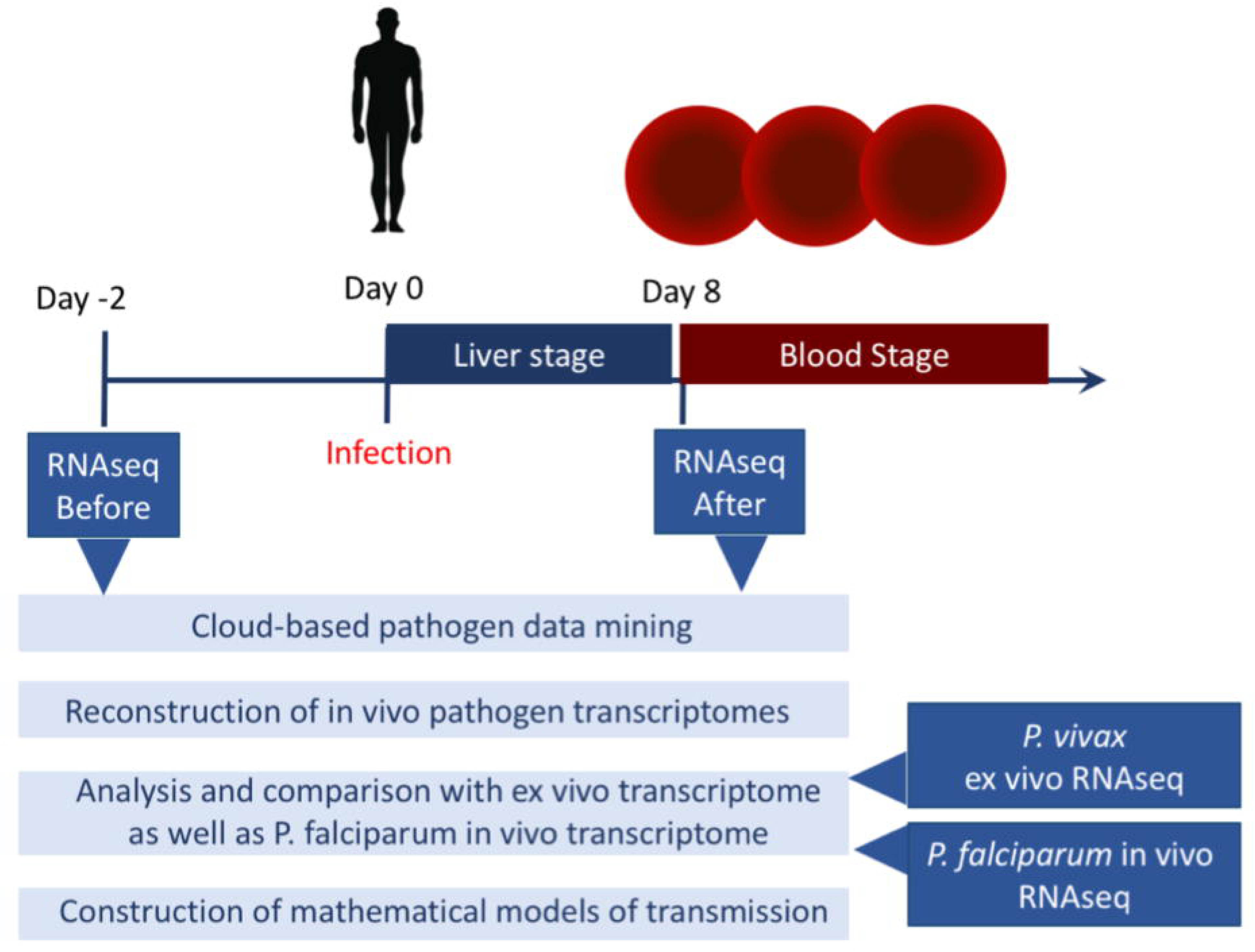
Study design and protocols. We have used two sets of RNAseq raw reads data pre and post sporozoite challenge from Rojas-Peña, et al. The post challenge data are inferred as the first blood stage cycle sequencing data. The early transcriptome signature is compared with publicly available *in vivo P. falciparum* and *ex vivo P. vivax* data to cross-validate the gametocyte signature in the early *in vivo P. vivax* infection.

## Material and methods

### Mining parasite data from infected human tissues

We used the blood transcriptome data sets deposited in Gene Expression Omnibus (GEO) under accession numbers GSE67184, GSE61252 associated with the *in vivo P. vivax* sporozoite challenge [11] and *ex vivo P. vivax* asexual stage culture [12] respectively. We also use the *in vivo P. falciparum* infection genomic reads [13] deposited in DNA Data Bank of Japan (DDBJ) under accession number DRA000949 to compare the transcript abundances with the above datasets.

PathoScope 2.0 [14] framework is used to quantify proportions of reads from individual species present in sequencing data from samples from environmental or clinical sources. A spot Elastic Computing Cloud (EC2) instance r3.4xlarge (Virtual CPUs – 16, Memory (GB) – 122, Storage (SSD GB) – 320)) was deployed at pricing of $0.13/hour. All the computational storage was synced with Amazon Simple Storage Service (Amazon S3), which automatically scales according to the current usage requirements. This facility gave us a cost effective ($0.03 per GB) advantage over the fixed storage on the local computing cluster. We used the Patholib module along with National Center for Biotechnology Information (NCBI) vast nucleotide database to create filter genomes containing host (human), microbes (virus, bacteria), artificially added sequence (PhiX Control v3, Illumina) and target genome library containing *P. vivax Sal-1* sequences using their respective taxonomic identifiers. PathoMap module is used to align the reads to target library using the Bowtie2 algorithm [15] and then filters reads that aligned to the filtered genomes. PathoReport was used to annotate the sequences.

The Tuxedo suite [16] of programs (Bowtie2, TopHat2, and Cufflinks) were used to process and analyze the data. Reference genomes of Human (*GHRc37*) from Ensembl human genome database and *P. vivax Sal-1* from PlasmoDB—a Plasmodium genome resource. Bowtie2 [15] was used to build indexes of the reference genomes. RNASeq reads from each sample were aligned to the *P. vivax Sal-1* genome using TopHat2 (v. 1.4.1) [17]. A maximum of one mismatch per read was allowed. The mapped reads from TopHat were used to assemble known transcripts from the reference, and their abundance FPKM values were calculated for all genes (FPKM: fragments per kilobase of exon per million fragments mapped) using Cufflinks.

### Gene expression level estimation with Poisson Modelling (PM)

Poisson distribution has been widely used to estimate the background level of gene expression [18-20]. In this work, we used Poisson distribution to model the background expression level (x) for each patient.

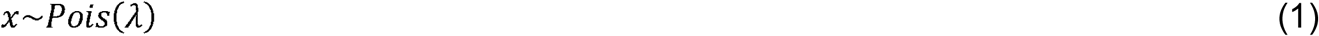

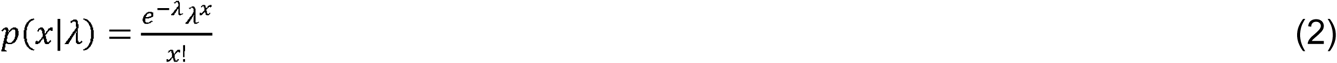

It is well known that the unbiased estimator of *λ* is the mean value of *x*, which can be calculated from maximum likelihood estimation.

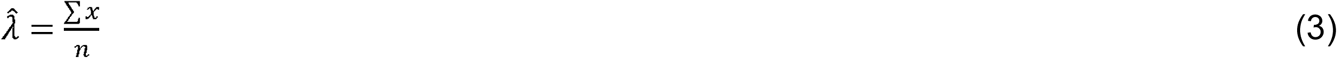

where ∑ *x* is the sum of gene expression level of specific patient or gene; *n* is the number of genes considered. Finally, we can compare the expression levels between different patients or genes by using the mean value of estimated distribution.

### Mathematical modelling of *P. vivax* transmission

The sexual stage specific genes are defined by using the 7 stages RNAseq data [21]. The stage specific RNAseq dataset is from Illumina-based sequencing of *P. falciparum* 3D7 mRNA from gametocyte stage II and gametocyte stage V), and ookinete. The dataset has also four time points of asexual stages representing ring, early trophozoite, late trophozoite, and schizont. The orthologs of *P. vivax* and *P. falciparum* were mapped with OrthoMCL data [22]. Sexual stage specific genes are required to have 20 or more fold expression level FPKM differences in the sexual stage (gametocytes, ookinete) vs the time points in the blood stages. The expression differences between asexual and sexual stages were analyzed with Fisher‒s Exact tests, and the P values (<0.001) were adjusted by multiple hypothesis correction with Benjamini-Hochberg method.

We created two mathematical models to represent the population-level spread of transmission among humans and mosquitoes for *P. falciparum* and *P. vivax* malaria. We do this to allow comparisons between the two malaria diseases, in order to assess which differences between the two have the most influence in producing the current epidemiological profile of the two diseases. This can inform us whether our genomics research results are an important aspect of *P. vivax* spread within populations. We categorize humans and mosquitoes into compartments based on their infection status, such as Susceptible, Exposed, Asymptomatic and Infected (see Supplementary Text S1), with additional compartments in the *P. vivax* model to account for the potential for relapses. For each model we calculate *R*_0_, the basic reproductive number of the disease. This is a commonly used, fundamental metric of disease transmission potential defined as the number of people one infected person is able to infect in a susceptible population. If *R*_0_ >1 then the disease is likely to take off and spread widely throughout the population. As the models for *P. vivax* and *P. falciparum* contain many similar components, we assess the relative *R*_0_ for the diseases, i.e. we divide all values of *R*_0_ for *P. vivax* by the value of *R*_0_ for *P. falciparum*. The model structure and resultant calculation of relative *R*_0_ allows us to easily make comparisons between the two diseases as well as ignore potential error in parameter values for those parameters which are shared between the two models.

In order to assess the impact of early transmission in humans on disease spread compared to other differences between *P. falciparum* and *P. vivax*, we perform a sensitivity analysis of *R*_0_ for *P. vivax*. For each parameter, we vary its value and calculate the new value of *R*_0_ to determine the effect of each parameter. We introduce parameter ε to represent the reduction in the length of the incubation period for *P. vivax* in comparison to *P. falciparum*; thus ε varies from 0 to 7 days to indicate a reduction from 14 to 7 days in the incubation period. That is, the larger ε is, the bigger a difference between *P. falciparum* and *P. vivax,* indicating earlier transmission for the latter disease. The parameter *p* represents the proportion of humans that are symptomatic in the *P. falciparum* model, and thus P varies from 0 to 1. In comparison, *k*_3_*P* indicates the proportion of symptomatic hosts in the *P. vivax model,* therefore, by focusing on *k*_3_ between 0 and 1, there are more asymptomatic cases for *P. vivax* than for *P. falciparum*. All other parameters are varied by 10% to create a range from 90% to 110% of the baseline value of each parameter. The more *R*_0_ changes when a parameter is varied, the more influence that parameter has on *R*_0_. In this way we can compare how much effect reducing the length of the intrinsic incubation period has on disease spread versus the role of asymptomatic spread or relapses.

Full details of the models created and the parameter values chosen as base values are presented in S1 Text.

## Results

### Using cloud-based computational pipelines to mine parasite derived transcript

To understand *P. vivax in vivo* pathogenesis, we first utilized a set of publicly available NGS raw data from Rojas-Pina et al. [11] that examined human immune responses against malaria. We performed computational analysis to extract the low levels of parasite signals from the raw sequencing data. The study by Rojas-Pina et al. performed sporozoite challenge on 12 volunteers with a single source of *P. vivax*, and generated whole blood RNAseq before and after the challenge. The post-infection RNAseq was produced on day 8-9 (diagnosis day), i.e., the first blood stage cycle after the liver stage infection which usually lasts for about 6-7 days [7, 8]. Due to the very low levels of parasitemia at this time point, we first used a cloud-based data mining pipeline to obtain pathogen sequences (*P. vivax*) in order to investigate the feasibility of our project. We deployed the program PathoScope 2.0 in the Amazon Elastic Compute Cloud (Amazon EC2: aws.amazon.com/ec2), due to the computational scalability that can be achieved within a few minutes. We mapped the entire set of raw sequencing reads to the NCBI NR(Non-Redundant) reference sequences and set *P. vivax* reference (Sal I) as targets. We have also used other pathogens such as viruses and bacteria as non-targets to increase the search specificity. From a total of 12 pairs of pre and post infection RNAseq raw sequencing reads data sets, we successfully detected *P. vivax* sequences from 1,000 to almost 50,000 reads in post-infection samples (Fig 2A). In contrast, none of the pre-infection samples gave a significant amount of reads (> 10). From this analysis, we concluded that we could precisely recover up to 50,000 P. vivax transcripts derived sequencing reads at this very early stage asexual replication.

**Fig 2.**
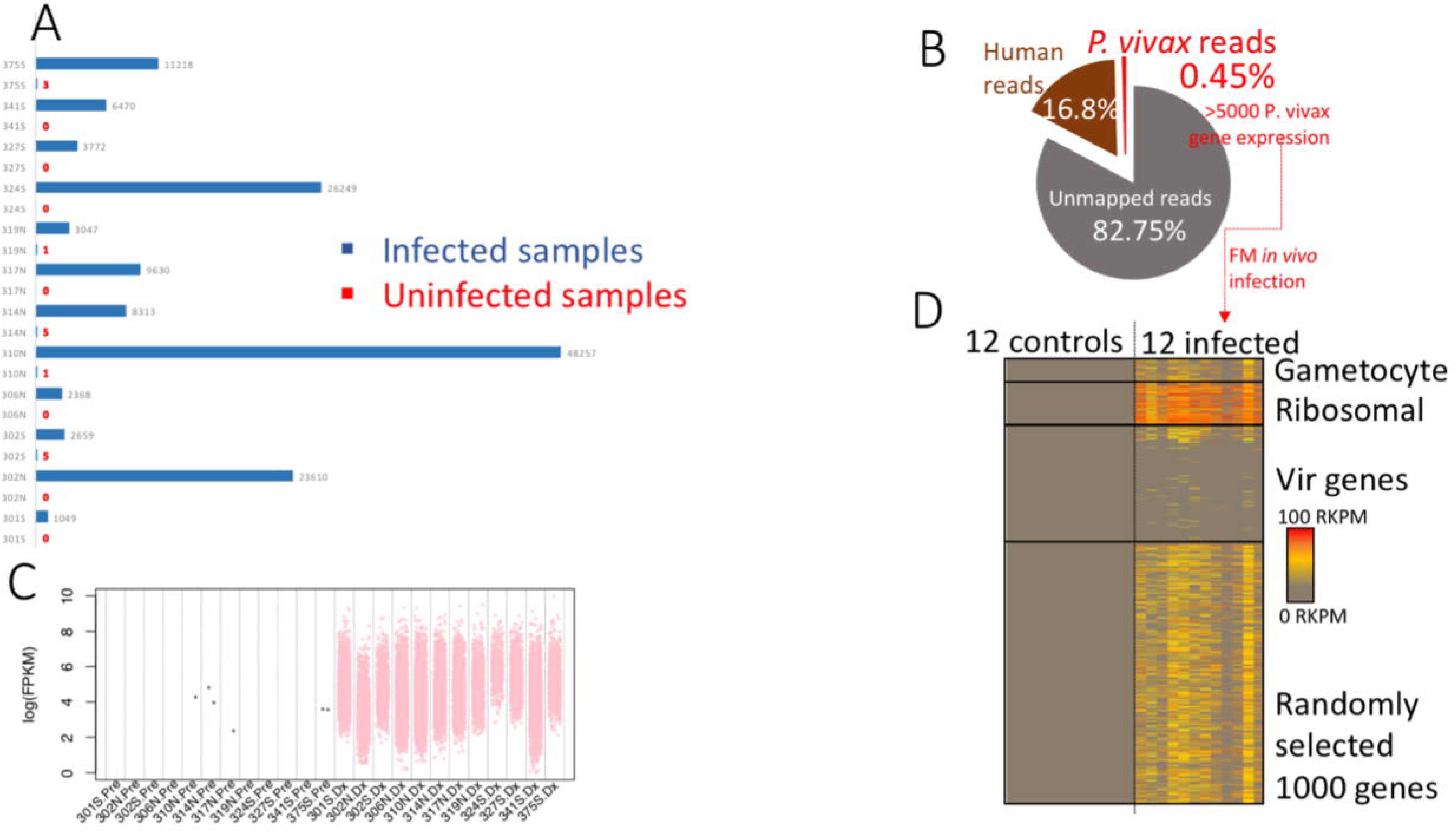
Recovering the earliest *in vivo* P. *vivax* blood stage transcriptome. Uninfected and post-infection blood samples were derived from the same individual. A total of 12 paired individual genomics data were analysed. **A**. Cloud-based sequence mining revealed that only the post-infection RNAseq raw data set contains parasite sequences in all patients. Patient identifiers are from the publication by Rojas et al. The *P. vivax* reads number is generated with stringent criteria and reflects conservative estimation. **B**. On average, less than 0.5% of total signal is derived from *P. vivax.* The mapped data of total reads and percentage of alignment in individual patient samples are listed in Table 1. C. The log(FPKM) distribution of all patients. FPKM represents fragments per kilobase of exon per million fragments mapped. Pre represents uninfected, while Dx means infected. Only the genes with FPKM **>** 0 are plotted here. **D.** RNAseq recovered parasite transcriptome in infected samples. Genes expressed in at least two patients are plotted.

**Table 1.** Patient specific information from literature and RNAseq data analysis.

| Patient Number | SRR [10] | Parasite Density on Pre-patent Day (Parasites/μL) [22] | Patient Location [10] | Total Reads | % reads aligned to Parasite Genome | % reads aligned to Human Genome |
| --- | --- | --- | --- | --- | --- | --- |
| 1 | SRR1925783 | 6 | Cali | 800452 | 0.32 | 17.52 |
| 2 | SRR1925785 | 10 | Cali | 1410398 | 0.24 | 16.92 |
| 3 | SRR1925803 | 20 | Buenaventura | 949274 | 0.09 | 16.1 |
| 4 | SRR1925797 | 25 | Buenaventura | 415781 | 0.19 | 19.52 |
| 5 | SRR1925781 | 34 | Cali | 725123 | 1.1 | 18.16 |
| 6 | SRR1925795 | 34 | Buenaventura | 446940 | 0.09 | 15.79 |
| 7 | SRR1925787 | 38 | Cali | 1055063 | 0.1 | 13.43 |
| 8 | SRR1925799 | 55 | Buenaventura | 587972 | 0.29 | 16.51 |
| 9 | SRR1925788 | 95 | Cali | 1570675 | 0.43 | 16.81 |
| 10 | SRR1925790 | 110 | Cali | 712547 | 1.13 | 18.48 |
| 11 | SRR1925798 | 216 | Buenaventura | 1238628 | 0.98 | 15.14 |
| 12 | SRR1925791 | 390 | Buenaventura | 711395 | 0.2 | 17.49 |

### Reconstruction of *P. vivax in vivo* transcriptome from very early blood stage infection

Next, we used the Tuxedo RNAseq pipeline [16] to reconstruct transcriptomes from the 12 post-infection samples, which is deployed in USF research computing cluster. We aligned the entire sequence data to *P. vivax Sal 1* and Human (*GHRc37*) reference genome and estimated the transcript abundances. The majority of the raw reads cannot be assigned to any references, primarily due to reads quality and possibly a small amount belonging to unknown genomes and the Phix179 control genome generally used during sequencing library construction. Reads originating from Phix were filtered prior to implementation of the Tuxedo RNAseq pipeline. An average of 16.8% of the reads can be mapped to human genome reference *GRCh37*. On average only 0.45% total reads on average mapped to the *P. vivax* reference genome (Fig 2B). From the 12 post-infection RNAseq pathogen transcriptomes, we can detect over 95% of the 5625 total protein coding genes expressed at > 20 FPKM (fragments per kilobase of exon per million fragments mapped). For each patient, we can identify from 9% to over 50% of the total protein-coding *P. vivax* genes are expressed at > 20 FPKM (S1A Fig).

To confidently identify parasite derived RNAseq signal from infected host tissues, we developed a Poisson Modelling (PM) method to characterize positive pathogen signals above the background. We modelled background signal with a Poisson distribution and estimated the significance of detected parasite transcriptional levels with a Maximum Likelihood method (S1 and S2 Table). To cross-validate our PM method, we have independently built our statistical model based on Negative Binomial Model (NBM) and obtained very similar results with only 0-2 genes expression in the control. Subsequently, we performed PM at two levels (S2A Fig). First, we used PM to evaluate patient level infection signals of before and after infection, taking the entire transcriptomes into account. Second, we used a gene-by-gene PM evaluation approach to identify the significantly expressed parasite genes in mixed sequencing results from human tissues (S2B Fig).

To understand the molecular patterns associated with *P. vivax* parasitemia, we designed a computational method to search for parasitemia associated genes. First, we grouped the patients into low (<=25 ul), medium (34-55/ul) and high parasitemia (95-300/ul) groups, based on the reported levels of parasitemia on pre-patent day, i.e. a range of 11 - 13 days [23], a few days later than the RNAseq samples were collected. We recognize that, in reality, all the patients have very few parasites during this early stage of infection, and the categories are primarily for statistical analysis. Then we performed a Non parametric statistical analysis (Wilcoxon test with p values adjusted with multiple hypothesis testing correction) to search for transcripts that are positively and significantly associated with the levels of parasitemia. In the top 20 *in vivo* parasitemia associated genes (p < 0.05), we identified genes with peak expressions in different asexual stages such as ring, trophozoite and schizont. We clearly identify a gametocyte expression signature at this early stage of *in vivo* infection in the top ranking markers.

### *P. vivax* parasitemia is associated with gametocytogenesis

To understand the extent of gametocytogenesis gene expression and its relationship with parasite abundance, we search for how many known gametocyte specific markers are expressed. We first defined a set of 280 gametocyte specific genes by using *P. falciparum* orthologous gene expression specificity (details in Materials and Methods). We discovered that between 8% to 60% all sexual stage specific genes are expressed in this early blood stage (S1B Fig). To further investigate the gametocytogenesis transcription pattern, we identified 48 gametocyte related genes from the patient infected transcriptomes. We were able to identify stage specific gametocyte markers with early markers [21] such as tubulin-specific chaperone PVX_081315 and Pvs16 PVX_000930. We also found late markers such as PVX_116610, indicating that there might be mixed stages of gametocyte obligation at this early blood stage. Furthermore, we have found gender specific markers, such as female markers like PVX_093600; as well as male markers such as PVX_116610 (S3 Table), indicating that there are mixed genders of gametocytes developed from a single source of sporozoite challenge. The transcriptional factor PvAP2-G is considered a master regulator and a specific marker for early gametocyte production in malaria parasites [24]. We are able to clearly identity transcripts of PvAP2-G in 5 patients in the early blood stage (Fig 3A).

**Fig 3.**
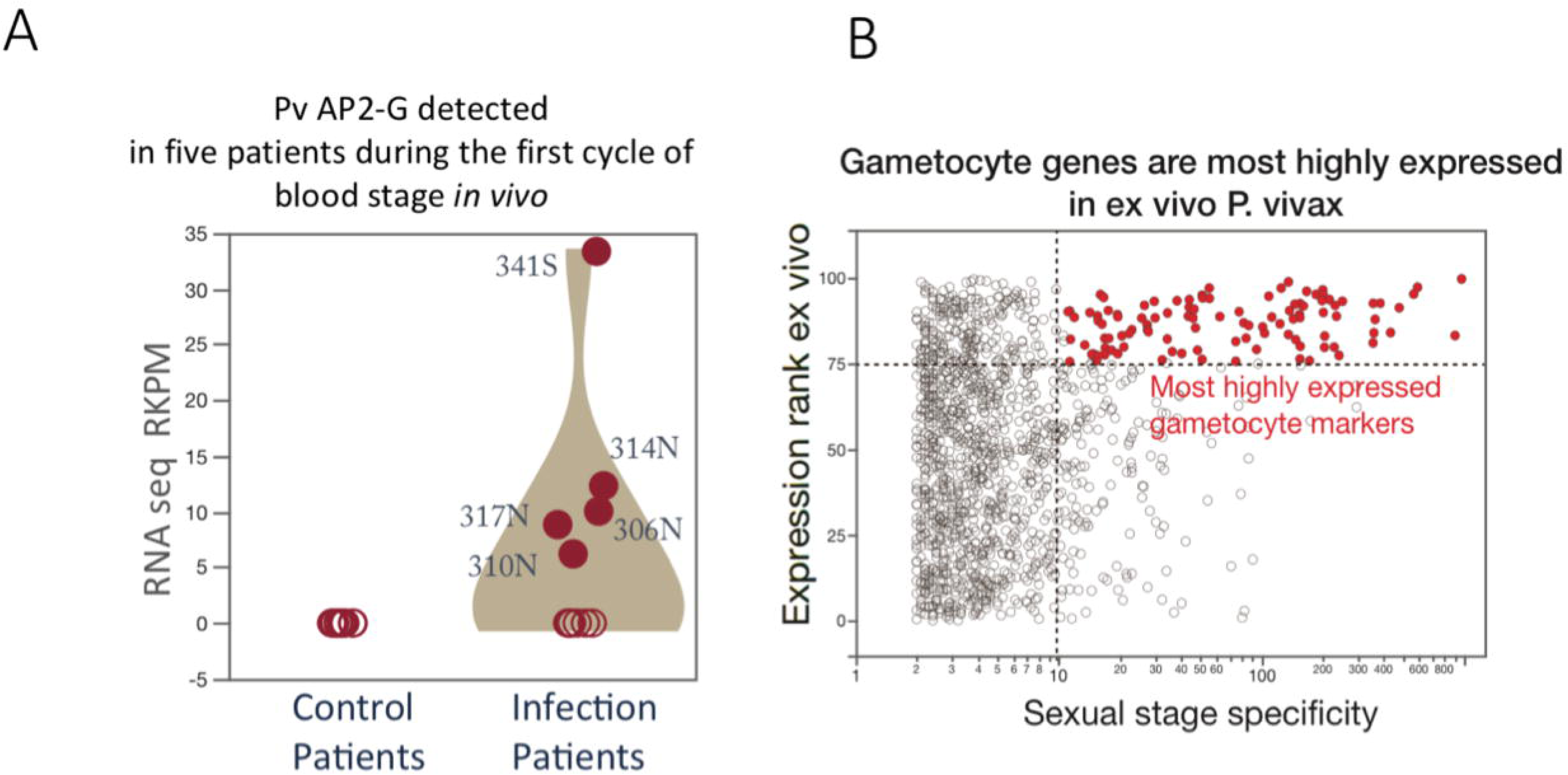
Discovering gametocyte signatures from early P. *vivax in vivo* RNAseq. **A.** Five patients showed expression of PvAP2-G, a master regulator of Plasmodium gametocyte production. **B.** Gametocyte specific genes are the most highly expressed genes in ex vivo P. *vivax* RNAseq transcriptomes. The *x* axis refers to the ratio of FPKM levels for sexual to asexual stages gene expressions. The top quartile of most highly expressed genes (Normalized rank score >=75) in the ex vivo data consisted of more than 40% of gametocyte specific genes.

To validate our findings of gametocytogenesis expression signature *in vivo*, we analyzed an independently generated, publicly available data set of *ex vivo* RNAseq data from pooled infected patient blood [12]. To search for relationships between expression levels and gametocyte production, we classified the *ex vivo P. vivax* expression based on two transcriptome features in orthologous genes of *P. falciparum*, namely, 1) Level of expression and 2) Specificity of gametocyte stage expression. We first computed the average FPKM for each gene and converted the values into a rank score from 0 to 100, with 100 representing the highest relative expression levels. Then we analyzed the levels of gametocyte expression specificity by calculating the ratio of sexual stage FPKM vs. asexual stage FPKM, the higher values indicate higher levels of sexual stage expression specificity. We found that the gametocyte specific genes in fact are the highest expressed genes in the *ex vivo* data (Fig 3B). The top quartile of most highly expressed genes in the ex vivo data consists of more than 40% of gametocyte specific genes. The *ex vivo* data has even stronger gametocyte expression pattern than that of the early *in vivo* data (S3A, B Fig). The *ex vivo* enhanced gametocyte induction could be due to the abiotic stress of the culture conditions (S3A, B Fig). By analyzing the precise peak expression time in the *ex vivo* expression data set, we found that gametocyte specific genes are mostly expressed in late schizont/early ring stage, despite the fact that these stages have the lowest number peak expression genes (S4A, B Fig). *P. falciparum* and *P. vivax* appear to share a pattern in which commitment to gametocyte development occurs in the schizont stage[25]. The *ex vivo* analysis strongly supports our *in vivo* analysis, that *P. vivax* parasitemia is associated with commitment to gametocytemia.

We next compared our *in vivo P. vivax* analysis with that of *P. falciparum*. Similar to *P. vivax* analysis, we have identified the top 20 transcripts associated with parasitemia from 120 whole blood samples of *P. falciparum* infected patients (data deposited in the publication by Yamagishi, et al.,) (Fig 4A, B). We defined these markers by searching for the gene expression levels that are most strongly associated in Spearman correlations with the levels of *P. falciparum* parasitemia among over 5000 unique transcripts. We found that in contrast to *P. vivax* parasitemia markers, *P. falciparum* parasitemia driven genes have peak expression only in the merozoite/early ring stages and many of them are associated with protein export as PEXEL containing proteins. None of the top *P. falciparum* markers are gametocyte related in terms of peak expression pattern. Therefore, we conclude that the two malaria parasites *in vivo* pathogenesis show distinct patterns. *P. falciparum* parasitemia is likely to be associated with asexual cycle protein export and host red blood cell remodelling; whereas *P. vivax* shows clear gametocyte expression signatures from the first blood stage cycle.

**Fig 4.**
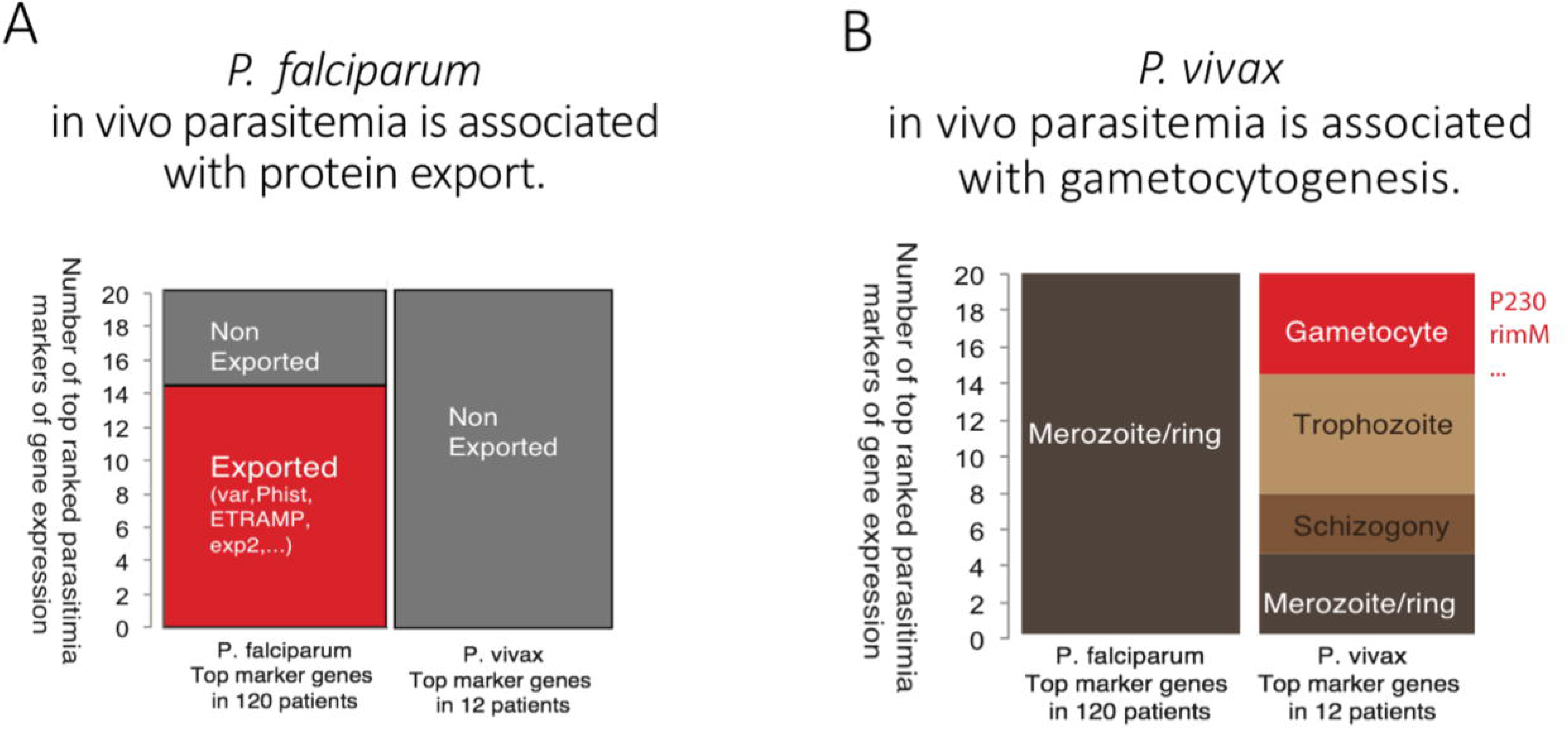
Comparison of *P. falciparum* and *P. vivax* in vivo transcriptomes. Top ranked markers that correlated with the levels of parasitemia are used for plotting. The top ranked parasitemia markers in *P. falciparum* are derived from 120 patients’ *in vivo* infection data. And the top ranked parasitemia markers in *P. vivax* are from 12 *in vivo* early infection data. **A**. Exported protein proportions in *P. falciparum* and *P. vivax.* Exported proteins are defined as PlamsoDBv27 PEXEL containing proteins; and they are likely be involved in host cell remodelling. **B**. Life cycle peak expression markers in *P. falciparum* and *P. vivax.* The peak expression patterns are assigned with all differentially expressed genes in 7 stages when there are more than 2-fold difference between stages.

### Mathematical modelling shows unique *P. vivax* transmission pattern

We performed a sensitivity analysis of the effect of different components of *P. vivax* disease spread (Fig 5). The most influential parameter on *R*_0_, the basic reproductive number of the disease, is *k*_2_, which determines the proportion of human hosts that recover with hypnozoites, and hence the possibility of relapse. It causes *R*_0_ to vary from less than 2.1 to over 2.5 times the values for *P. falciparum* (set to 1, when *p*= 1, where P is the proportion of *P. falciparum hosts that are symptomatic*). The second most influential parameter on changes in *R*_0_ is ε, the reduction in the length of the incubation period. When there is no difference between *P. falciparum* and *P. vivax*, that is ε is 0 days, *R*_0_ is lowered to less than 2.2. However, when the incubation period is shortened for *P. vivax* by *ε* = 7 days, as we expect from our experimental results, *R*_0_ is 2.4. Therefore, if the reduction in incubation time is not considered, mathematical models could miscalculate *R*_0_, underestimating it by approximately 11%. However, this assumes that reducing the time to potential transmission does not have any other impact on the disease characteristics. There could be a trade-off between the speed of gametocyte production and the efficiency of those gametocytes in transmitting the disease, and if so this would mitigate against the reduction in incubation length [26]. Parameters *v* and *η*, the rate of relapse and the rate of hypnozoite death in the liver respectively, are also influential in determining the value of *R*_0_, as are the parameters related to proportion of hosts that show symptoms, *p* and *k*_3_. Introducing asymptomatic hosts simultaneously to both *P. falciparum* and *P. vivax* (i.e. changing P) reduces the relative value of *R*_0_ for *P. vivax* because it has a larger impact on *P. falciparum*. However, *R*_0_ is more sensitive to parameter *k*_3_ than *p*, indicating that it is necessary to understand the likelihood of asymptomatic cases in *P. vivax* compared to *P. falciparum* to accurately predict differences in disease spread. The influence of these parameters highlights the importance of understanding the role of the asymptomatic stage correctly.

**Fig 5.**
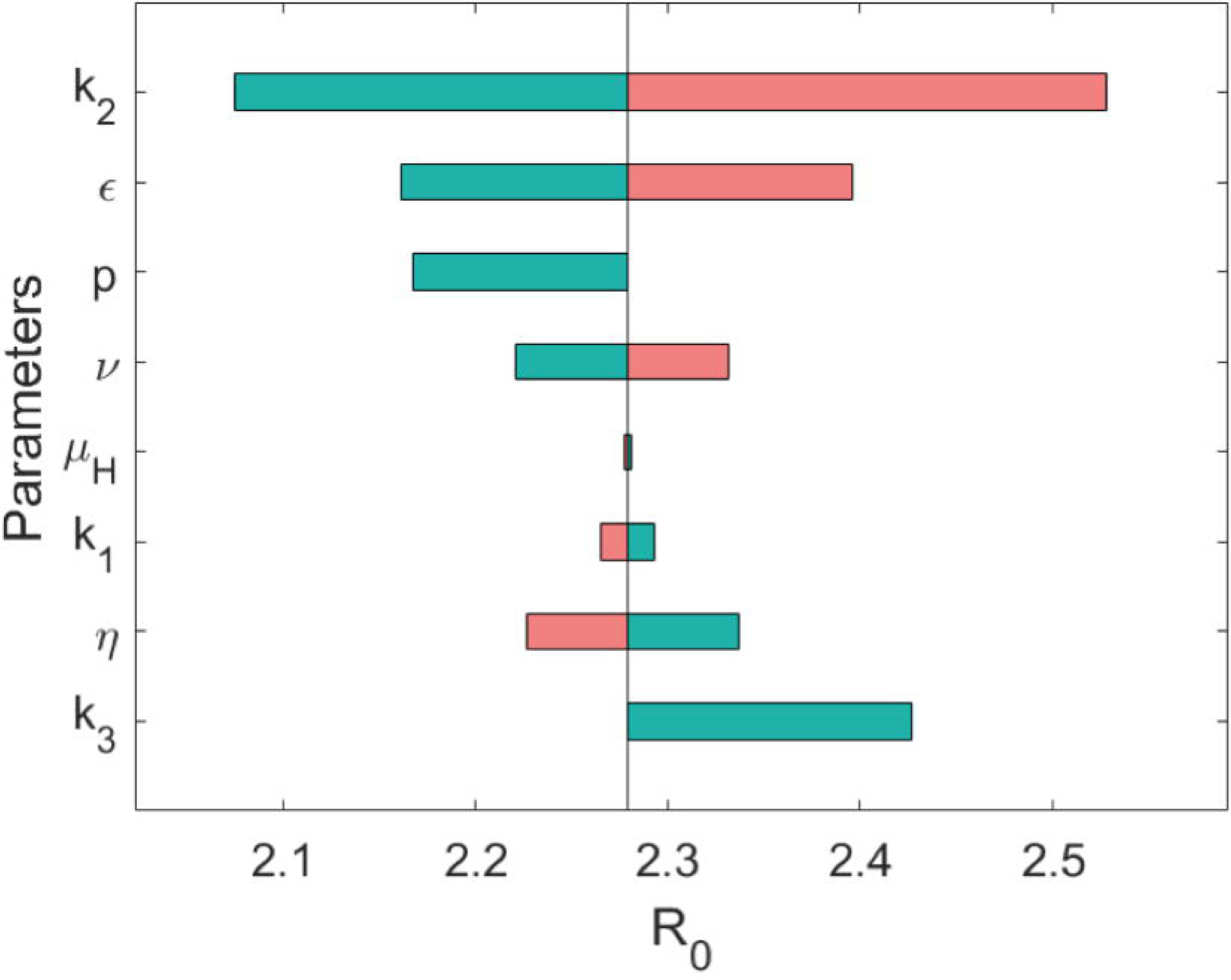
Mathematical model of P. *vivax* exploring the effect of reduced incubation period on spread of disease. A sensitivity analysis is performed on *R*_0_ for *P. vivax* (relative to *P. falciparum*). Green indicates when the parameter has been lowered from its baseline value, pink indicates higher than baseline (therefore *R*_*0*_ is positively correlated with the first four parameters and negatively correlated with the last four parameters). Parameters *ε*, *p* and *k*_3_ are varied between 0 ‒ 7, 0 ‒ 1, and 0 ‒ 1 respectively, all other parameters are varied by 10%. Parameters are: proportion of hosts that develop hypnozoites (*k*_2_), reduction in incubation time (*ε*), proportion of hosts developing symptoms in *P. falciparum* (*p*), rate of relapsing (*v*), host death rate (*µ_H_*), proportional rate of disease-induced death for *P. vivax* (*k*_1_), rate of hypnozoite death in liver (*ƞ*) and proportion of hosts developing symptoms in *P. vivax* relative to *P. falciparum* (*k*_3_). Parameter values are in S1 Text.

We further explore the role of the reduction in the incubation period length in Fig S5, which shows the effect of not including relapses in the model and not accounting for asymptomatic hosts transmitting the infection in *P. vivax*. When we model the asymptomatic class as capable of transmitting infection but are unsure what proportion of hosts are in this category, our uncertainty in *R*_0_ is small (Fig S5A). On the other hand, when asymptomatic hosts for *P. vivax* exist but the existence of these asymptomatic cases is unknown and hence not modelled as capable of spreading disease, there is a drastic underestimation of *R*_0_, the potential for spread, of *P. vivax* (Fig S5B). In fact, if more than 40% more infectious hosts are asymptomatic compared to *P. falciparum*, the estimate of *R*_0_ for *P. vivax* would be less than for *P. falciparum* when in reality it is approximately 2.5 times larger. Similarly, when the model does not account for relapses, the estimate for *R*_0_ is halved (Fig S5B).

## Discussion

Our study has uncovered the earliest possible *in vivo* infection data of blood stage *P. vivax*, a parasite that cannot be cultured in the laboratory. Our study is thus an example for infectious disease researchers on how to use large raw sequencing data to investigate previously intractable pathogenesis-related features. We used a cloud-based mining method as part of our study. This approach does not require local High Performance Computing (HPC) facilities and can accommodate high volumes of data analysis within short time frames. Infectious disease scientists could use similar approaches in resource-limited research settings. As the publicly available genomic data grow in complexity and volume every day, more efficient and more precise analytical tools are needed for future studies.

With malaria eradication always in the spotlight of the scientific and public health community, there is an urgent need to understand the unique biological and physiopathological features of *P. vivax*. If, as our data suggest, *P. vivax* transmission to mosquitoes is plausible at the very first blood stage cycle immediately after liver stage development, this would represent a major hurdle towards targeting *P. vivax* reservoirs. Due to ethical and practical limitations to obtain experimentally infected *P. falciparum* data *in vivo*, our study used *P. falciparum* data without defined infection age. Nevertheless, the major differences we have discovered between *P. vivax* and *P. falciparum*, in terms of *in vivo* gene expression, suggest that *P. vivax* begins gametocyte production immediately upon entering the blood, whereas more research is needed for early gametocyte production in *P. falciparum*.

Early stage I gametocytes of *P. falciparum* can be initially in peripheral blood and are microscopically indistinguishable from early rings [27, 28]. Yet, we did not find strong transmission expression signatures. It stands to reason that the 1-2 weeks of bone marrow sequestration that *P. falciparum* needs in order to achieve a fully transmissible stage V truly represents an advantage for *P. vivax* transmission over *P. falciparum*. Further, it has been described [29] that even very few gametocytes in circulation, as inferred from our study in *P. vivax*, can effectively mount an infection in the mosquito host. For asymptomatic infections, although the evidence is mixed and it has been suggested that the proportion of symptomatic and asymptomatic clinical forms is roughly similar for both species, when studied in native Amazonian populations [30], others have reported that the relative proportion of submicroscopic *P. vivax* is significantly higher than that of *P. falciparum* [31, 32]. Taking into account that over 89% of *P. vivax* submicroscopic infections are said to be asymptomatic [33], the balance in terms of better asymptomatic transmissibility falls on the side of *P. vivax*. Altogether, these evidence suggests that the differences we have discovered between *P. vivax* and *P. falciparum*, in terms of *in vivo* gene expression, suggests that *P. vivax* has the ability to spread quickly to multiple hosts before the onset of symptomatic phenotypes.

Our mathematical model, which accounts for *P. vivax* relapses, re-enforces the idea that *P. vivax* will be more difficult to eliminate. Hence, our results confirm the idea, held widely, that *P. vivax* will be the last parasite standing before the goal of malaria eradication is to be achieved [3]. Our model serves as a framework for further simulation and a better understanding of *P. vivax* population dynamics. It can also be adapted to account for the potential evolutionary consequences of reducing the length of the incubation period. A shorter incubation period could indicate lower production of efficient gametocytes, therefore the probability of successful transmission from an infected human to mosquito could be reduced. This could be achieved by introducing a trade-off function between these two parameters in the model. However, the form of this trade-off function is not clear and would need to be investigated experimentally. The potential reduction in incubation period, and hence early transmission, has a substantial impact on disease spread, dependent on this evolutionary trade-off. Without including shorter incubation periods, models may underestimate the work required to reduce transmission of *P. vivax* within a population. Our model also highlights the importance of relapses and asymptomatic carriers, with relapses the most influential factor in leading to increases in disease spread. And yet, relapses are poorly understood with no consensus on what causes relapses to occur or on their frequency. Further, the importance of asymptomatic carriers is interesting as *P. vivax* has long been associated with milder disease symptoms and many patients could be asymptomatic [34]. Our model highlights the importance of including the asymptomatic stage within models, even if the exact proportion of hosts that will not show symptoms is unknown.

Overall, the unique transmission of *P. vivax* leads to a much higher likelihood of disease spread compared to *P. falciparum* in similar settings. Our work highlights the challenge of *P. vivax* eradication and provides evidence for the need for more thorough and earlier transmission intervention measures. Controlled transcriptomic studies comparing *P. falciparum* and *P. vivax* gametocyte gene expression in oocysts and sporozoites are needed in order to understand how soon sexual commitment is decided in the *P. vivax* complex life cycle. Since *P. vivax* commits to gametocytogenesis early in the blood stage rationally designing a treatment or vaccine targeting the early blood stage will reduce transmission rates.

## Acknowledgement

We would like to thank Justin Gibbons, Alison Roth and John H Adams for constructive discussions.

